# Matrix Linear Models for connecting metabolite composition to individual characteristics

**DOI:** 10.1101/2023.12.19.572450

**Authors:** Gregory Farage, Chenhao Zhao, Hyo Young Choi, Timothy J. Garrett, Katerina Kechris, Marshall B. Elam, Śaunak Sen

**Author notes:** Use footnote for providing further information about author (webpage, alternative address)—*not* for acknowledging funding agencies (Gregory Farage).

## Abstract

High-throughput metabolomics data provide a detailed molecular window into biological processes. We consider the problem of assessing how the association of metabolite levels with individual (sample) characteristics such as sex or treatment may depend on metabolite characteristics such as pathway. Typically this is one in a two-step process: In the first step we assess the association of each metabolite with individual characteristics. In the second step an enrichment analysis is performed by metabolite characteristics among significant associations. We combine the two steps using a bilinear model based on the matrix linear model (MLM) framework we have previously developed for high-throughput genetic screens. Our framework can estimate relationships in metabolites sharing known characteristics, whether categorical (such as type of lipid or pathway) or numerical (such as number of double bonds in triglycerides). We demonstrate how MLM offers flexibility and interpretability by applying our method to three metabolomic studies. We show that our approach can separate the contribution of the overlapping triglycerides characteristics, such as the number of double bonds and the number of carbon atoms. The proposed method have been implemented in the open-source Julia package, `MatrixLM`. Data analysis scripts with example data analyses are also available.

## 1 Introduction

High-throughput metabolomics provides a detailed molecular window into biological processes involved with low-molecular-weight compounds referred to as metabolites. Metabolomics analysis deals with quantitative and qualitative assessments of metabolites, such as lipids, amino acids, nucleic acids, peptides, and steroids. Numerous studies have already shown the tremendous potential of metabolomics analysis in research fields as diverse as toxicological mechanisms, disease processes, and drug discovery (Zhang *et al*., 2015). For instance, metabolomics permits distinguishing between normal and pathological pathways, helping diagnose disease, and predicting prognosis (Aderemi *et al*., 2021). Various data mining and statistical methods can be used in metabolomics studies depending on the experimental context (Xi *et al*., 2014; Chen *et al*., 2022). The metabolome can be used as a predictor for a trait or phenotype, or it can be the response variable of interest as well. In this work, we consider the case when the metabolome is regarded as the response whose variation we want to predict based on metabolite or individual characteristics.

The first step in analyzing the metabolome is usually a univariate analysis of each metabolite. For two-group data, fold change analysis, t-tests, and volcano plots are common approaches. When analyzing multi-group data, one-way ANOVA (analysis of variance) and correlation analysis can be used. Multiple testing methods such as Bonferroni correction, Bonferroni-Holm correction, or Benjamini–Hochberg (also known as the false discovery rate, FDR) are used across metabolites to control the false positive rate. Often, the univariate analysis is followed by an enrichment analysis or similar to find patterns among the discoveries (Chen *et al*., 2022). The end goal is to identify metabolite characteristics (*e*.*g*., lipid class or subclass, carbon chain length,…) that have different levels among the groups. Put differently, the analysis aims to identify metabolite characteristics associated with the sample/individual characteristics (groups). We wish to answer questions such as: are unsaturated triglycerides (metabolite characteristic) associated with fish oil consumption (individual characteristic)?

There are two challenges when looking at patterns across metabolites that are associated with a sample characteristic of interest. The first challenge is that metabolites are often correlated, sometimes quite strongly, and usually, enrichment analyses ignore such correlations. The second issue is that some metabolite characteristics may not be categorical and may be quantitative, which enrichment analysis cannot handle. For example, triglycerides can be characterized by the number of carbon atoms and the number of double bonds they have.

Multivariate analysis techniques can be beneficial to address the challenge of correlated metabolites. Factor analysis, PCA (principal component analysis), PLS-DA (partial least squares discriminant analysis), and cluster analysis have been used in this context. A multivariate analysis may begin with an unsupervised technique (Huang *et al*., 2022; Chen et al., 2022), *e*.*g*. PCA, followed by a supervised classification model and identifying key metabolites through variable importance scores. PCA is often utilized as a preliminary analysis and quality control measure for metabolomics data to examine whether there are distinctions between groups and outlier points (Huang *et al*., 2022). Additionally, PCA can be used to assess whether quality control samples cluster together or if they show discrepancies which could suggest problems with the assay’s accuracy. Being an unsupervised analysis, PCA may be less readable and interpretable than the original variables. In addition, if caution is not taken when choosing the number of principal components, some information may be lost in comparison to using the initial set of features (Nyamundanda *et al*., 2010), as PCA aims to capture maximum variance amongst all variables. PLS-DA is a type of machine learning algorithm used for classification that attempts to reduce the number of variables by creating linear combinations and taking into account class labels. It tries to identify important factors while eliminating those that don’t contribute to predicting the label. However, it can be vulnerable to overfitting as it relies heavily on the initial training data to make accurate predictions (Gromski *et al*., 2015). While it can be useful for prediction, it does not directly help us analyze the metabolome as a response variable. Random Forests (RF) are helpful in finding nonlinear patterns between metabolites and outcomes in metabolomics (Chen *et al*., 2013). However, the RF feature selection process may not be effective if the variables have different scales of measurement or different numbers of categories (Gromski *et al*., 2015). RF does not also provide measures of statistical significance or p-values, only a ranked list of the most important metabolites (Chen *et al*., 2022). The other mentioned machine learning methods have similar limitations to RF in terms of providing variable importance measures but not offering statistical significance. In order to make an objective and comprehensive evaluation of each variable importance, it is a common strategy to use univariate analysis first, then combine the importance scores from multivariate models as screening criteria, and finally integrate them with variables selected by machine learning models such as those with FDR less than 0.05 or that are ranked high in the RF for potential biomarkers.

We propose using matrix linear models (MLMs), a family of bilinear models, to analyze correlated metabolomics data and aggregate signals from metabolites with similar characteristics (quantitative or categorical) and associate them with individual characteristics of interest. We developed the MLM framework (Liang *et al*., 2019) for studying associations in structured high-throughput data. This technique aggregates signals from both samples and omics features in a single model, and our efficient estimation algorithm exploits its structure to extract information. The MLM framework contrasts with the conventional metabolomics approaches, which analyze each metabolite independently and then look for patterns among those displaying similar associations. We have already applied this method to high-throughput genetic screens where we show that it is more powerful compared to analyzing features one at a time. It is related to LIMMA (Ritchie *et al*., 2015), which fits linear models for each metabolite separately, and then aggregates signals using an Empirical Bayes approach but does not explicitly use any metabolite characteristics or annotations. Our approach explicitly uses the metabolite characteristics in bilinear models. Below, we demonstrate its flexibility through three real-world metabolomic studies.

The paper is organized as follows. In Section 2, we introduce three distinct metabolomics studies that demonstrate the application of Matrix Linear Models (MLM) for analyzing metabolomic data. Section 3 provides the theoretical framework behind matrix linear models (MLM), detailing its computational strategies. We introduce the metabolite covariate matrix (“*Z* matrix”) that brings in information on the metabolites. In Section 4, we present the results from the application of MLM in the three distinct datasets. The paper concludes, in Section 5 with a summary of the key findings, implications for the broader metabolomics research field, and suggestions for future studies employing M.

## 2 Framing Metabolomic studies

In this section, we introduce three metabolomics studies that we revisited to illustrate the use of MLMs. Each study focuses on a different biological problem using metabolomics which we hope will give the reader an impression of the variety of problems that can be tackled with MLMs.

### 2.1 Statin-associated muscle symptoms study

Garrett et al. (2023) explored the metabolomic and lipidomic profiles in plasma samples from patients undergoing statin rechallenge, a part of their clinical care for statin-associated muscle symptoms (SAMS). While statins, widely known for their cholesterol-lowering capabilities and benefits in cardiovascular disorders, are the primary treatment context, our paper’s focal point diverges to a specific, incidental confounding factor: the impact of fish oil supplements.

Using a liquid chromatography-mass spectrometer (LC-MS), we analyzed samples from 98 patients, including those who had consumed fish oil supplements. We directed our analysis toward understanding how fish oil supplementation might influence triglyceride composition. In this study, 787 identified triglycerides were examined through matrix linear models to delineate this association.

Statins are effective and widely popular cholesterol-lowering agents with substantial benefits for preventing and treating cardiovascular disorders. However, about 10% of statin users cannot tolerate this drug (Bytyçi *et al*., 2022). The most common reasons for statin intolerance are usually related to muscle pain, *i*.*e*. SAMS. This common disorder causes poor adherence to statin therapy, which subsequently may alter their efficacy and increase the occurrence of cardiovascular events. With no validated biomarkers or tests to confirm SAMS, it remains particularly difficult to diagnose this side effect since it heavily relies on patients self-reporting their symptoms. The ability to identify patients at risk of SAMS would adapt the approach to treatment using lower-intensity statins or by lowering the doses of statins and combining them with non-statin LDL lowering therapy.

Although statins are the background of our study due to their role in the treatment regime of our participants, the main objective is to examine the effects of fish oil supplements on lipid profiles, especially triglycerides. Our analysis adds a unique dimension by analyzing fish oil supplementation and its potential implications in the lipidomic landscape of patients under statin treatment.

### 2.2 Pansteatitis Mozambique tilapia study

In an *in situ* study conducted at Loskop Dam, South Africa, Koelmel et al. (2019) used liquid chromatography coupled with high-resolution tandem mass spectrometry to analyze the plasma lipidome of 51 Mozambique tilapia *(Oreochromis mossambicus)*. They measured 590 distinct metabolites to find lipid markers differentiating healthy tilapia from those affected by pansteatitis. However, a key aspect of our research is extending this analysis to examine how the age of these fish influences the composition of various lipid groups.

The collection region witnessed mass mortality events involving both fish and Nile crocodiles *(Crocodylus niloticus)*, attributed to an unknown cause prompting the spread of pansteatitis, a debilitating inflammatory disease of adipose tissue leading to impaired mobility and death.

The emerging subfield of clinical lipidomics has already shown great promise in clinical science for biomarkers and mechanisms of disease (Meikle *et al*., 2021). However, analyzing lipid profile fluctuations in wildlife and environmental studies poses unique challenges since those profiles might be susceptible to natural changes across time and geography. Consequently, *in situ* studies offer opportunities to document baselines that contrast lipid profile changes due to naturally occurring phenomena versus the studied environmental change.

### 2.3 COPD-SPIROMICS study

Gillenwater et al. (2021) analyzed two large cohorts, COPDGene and SPIROMICS, to investigate whether there are sex-specific metabolomic differences between COPD (chronic obstructive pulmonary disease) and emphysema. The COPDGene dataset featured 999 metabolites and had 839 participants, while the SPIROMICS dataset contained 787 metabolites and had 446 participants. These studies have looked at metabolomic profiles using blood samples, specifically plasma, since they are the preferred option for biomarker discovery due to their non-invasive nature and easy accessibility. Accurately characterizing sex-specific molecular differences in COPD is crucial for personalized diagnostics and therapeutics. Gillenwater et al. (2021)’s definition of COPD case status was based on spirometric criteria indicating at least moderate airflow obstruction, which was defined by a post-bronchodilator Forced Expiratory Volume in one second to Forced Vital Capacity ratio (FEV1/FVC) of less than 0.50, coupled with an FEV1 percent predicted (FEV1pp) below 80%. The control subjects were identified by a FEV1/FVC ratio greater than 0.7 and a FEV1pp exceeding 80%.

The COPDgene and SPIROMICS plasma samples were profiled using the Metabolon (Durham, USA) Global Metabolomics Platform. The extracted samples were partitioned into four distinct fractions for comprehensive analysis. Two fractions were allocated for analysis using reverse-phase/ultrahigh performance liquid chromatography-tandem mass spectrometry (RP/UPLC-MS/MS) equipped with positive ion mode electrospray ionization (ESI). Another fraction was dedicated to RP/UPLC-MS/MS analysis but utilized negative ion mode ESI. The fourth fraction was reserved for analysis through hydrophilic interaction chromatography (HILIC)/UPLC-MS/MS, again employing negative ion mode ESI.

## 3 Methods

In this section we outline the statistical framework underlying matrix linear models, and the software implementation in the Julia programming language.

### 3.1 Matrix Linear Model

MLMs provide a framework for studying associations in high-throughput data when the response or output is a large matrix, and we have annotations on both the rows and columns of that data matrix. We developed this framework in the context of high-throughput genetic screens (Liang *et al*., 2019). Outline below is the application of our framework to metabolomics studies. The data structures involved are represented in Figure 1. Consider the metabolomics profile of all individuals as the data (response or output) matrix, we would have additional information (annotations) on the rows of that matrix (covariates corresponding to individuals) and the columns (attributes corresponding to each metabolite). It operates on a matrix of expression values, each column representing an omic feature relevant to the current study, here metabolites and each row corresponding to a sample. Our method may be viewed as a generalization of Ritchie *et al*. (2015) in that we use a bilinear model in both the response attributes and the individual attributes; the former uses a linear model in terms of the individual attributes for each metabolite.

**Figure 1:**
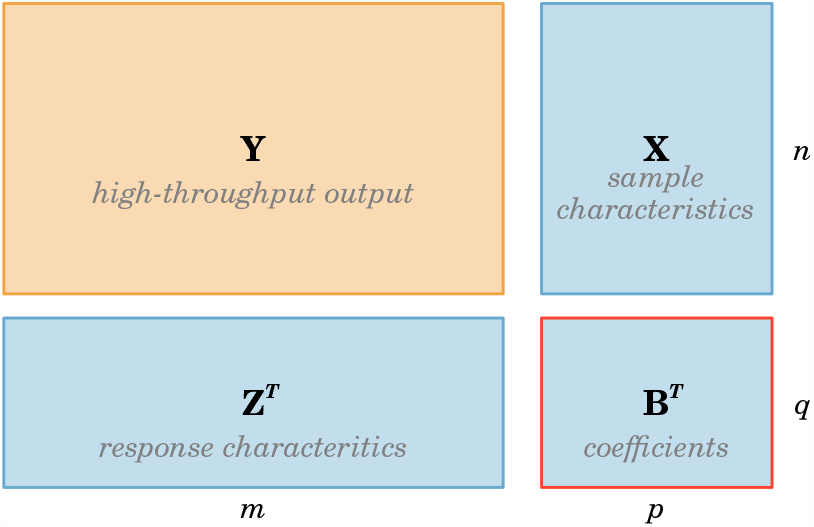
A visualization of the response (**Y** : *n × m*), covariates (**X** : *n × p*), measurement attributes (**Z** : *m × q*), and coefficients (**B** : *p × q*) matrices for a matrix linear model. The dimensions in the model correspond to *n* individuals, *m* measurements, *p* covariates, and *q* attributes; the matrix **B** is to be estimated.

Our bilinear model may be expressed in matrix form as

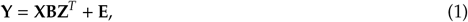

which is equivalent, in summation notation, to

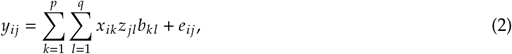

where we denote *a*_*ij*_ as an entry of the matrix **A**. The matrix **Y**_*n*_×_*m*_ is the output data arranged in *n* samples and *m* types of measurements (*e*.*g*., triglyceride levels); the matrix **X**_*n*_×_*p*_ consist of the covariates composed of *n* samples and *p* independent variables (*e*.*g*., phenotype status, whether they were taking fish oil); **Z**_*m*_^×^_*q*_ is the matrix of measurement attributes composed of *m* measurements and *q* attribute variables (*e*.*g*., information about triglycerides); the matrix **B**_*p*_^×^_*q*_ holds the coefficients that must be estimated (*i*.*e*., can be interpreted as interactions between **Z** and **X** covariates); the random error matrix **E** is assumed to have a mean zero, independent across individuals, but possibly correlated across outcomes, such that *V ar*(*vec* (**E**) ) = **Σ** ⊗ **I** where **Σ** is the residual covariance matrix.

When there is one metabolite (or **Y** has a single column) and there is no **Z** matrix (response attributes), then the model reduces to linear model for the one metabolite in terms of individual attributes (encoded in the matrix **X**).

We calculate the coefficients by employing least squares estimation, which means that we choose the matrix **B** in order to minimize the sum of squared residuals(3),

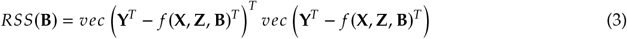

where *f* **X, Z, B** = **XBZ**^*T*^ . The solution can be viewed as a generalized estimation equations approach (Xiong *et al*., 2011) a closed-form solution.

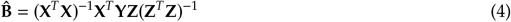

where the variance-covariance matrix of the estimated coefficient 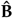 is defined as

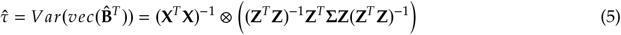

We can define a test statistic for our method that is analogous to the t-test statistic used to evaluate the coefficients from the univariate linear regression model,

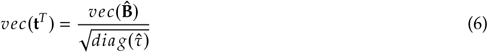

where **t** is the matrix of statistics of the estimated coefficients 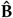, and 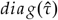 is the vector of the diagonal entries of the variance-covariance matrix of 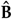. We provide a detailed derivation of the least squares estimate and variance in the appendix of this paper (Liang *et al*., 2019), where we have already used this approach to chemical genetic experiments and achieved positive outcomes, as evidenced by simulated data.

The **Z** matrix offers some leeway in designing feature interaction between sample features and metabolites groups. This approach allows us to identify patterns and detect associations that would not be possible if the data were split into subsets for pairwise comparison.

In our statistical analysis, we computed 95% confidence intervals, and to account for multiple testing, we employed the adaptive Benjamini–Hochberg method (Benjamini *et al*., 2006).

### 3.2 Software implementation in Julia

We have implemented MLMs in the `MatrixLM` package written in the Julia programming language. Users can specify how they want to model the individual covariates (*X*) and the response attributes (*Z*) using model formulas using the @formula macro. The package estimates the coefficient matrix and associated standard errors. Users can also obtain test statistics using permutation tests, predicted values, and simulate data. Some users may also be interested in a related Julia package MetabomomicsWorkbenchAPI that can be used to download data from the Metabolomics Workbench website using their application programming interface (API).

## 4 Data analysis

In our Matrix Linear Model (MLM) analysis, we define an “adjusted model” as one where multiple annotation variables are included alongside the primary variable of interest. For example, an adjusted model might incorporate variables such as carbon chain length and the number of double bonds. This approach allows us to assess the effect of the primary variable(s) while simultaneously accounting for the influence of these additional annotation characteristics. Conversely, an “unadjusted model” refers to a less complex MLM setup where only one annotation variable is included in addition to the primary variable of interest. In this scenario, the analysis focuses more directly on the relationship between the primary variable(s) and the outcome without the simultaneous consideration of multiple additional annotation variables.

### 4.1 Statin-associated muscle symptoms study

#### 4.1.1 Descriptive Characteristics

Patients who had been diagnosed with SAMS (Cases) were recruited from the Lipid Metabolism Clinics at the Memphis VA Medical Center, while Controls were recruited from the general patient population. Eligible Cases must have discontinued statins on two or more occasions due to muscle weakness or myalgia and be willing to undergo rechallenge with a statin medication; additionally, they had to meet certain criteria, including a myalgia clinical index score of 9-11 (based on the National Lipid Association definition of probable SAMS) or Naranjo probability score for adverse drug reactions. Exclusion criteria included advanced renal disease, active liver disease, advanced cirrhosis, clinically active autoimmune disease, and current use of one or more drugs contraindicated for statin use. Controls had to have been compliant with statin therapy for at least one year based on drug dispensing records and had confirmed the absence of SAMS symptoms during that time. Both Cases and Controls were studied while on statin therapy; the rechallenge period lasted up to 4 weeks or until myalgia symptoms developed in the Case group, after which blood samples were collected from all participants following an overnight fast. Some patients from both the case and control groups used fish oil supplements in addition to taking statin drugs. Table 1 displays the multivariate frequency distribution for both variables: group (case and control) and fish oil supplement status.

**Table 1:** Two-way contingency table for SAMS and fish oil supplement status.

| Supplement | Control | Cases | Total |
| --- | --- | --- | --- |
| Fish Oil | 39 | 26 | 65 |
| No Fish Oil | 15 | 18 | 33 |
| Total | 54 | 44 | 98 |

**Table 2:** Descriptive statistics of Mozambique tilapia samples.

| Characteristics | Diseased (N = 30) | Healthy (N = 14) | Total (N = 44) | P-value |
| --- | --- | --- | --- | --- |
| Sex, n(%) |  |  |  | 0.4 |
| Female | 15 (50) | 5 (36) | 20 (45) |  |
| Male | 15 (50) | 9 (64) | 24 (55) |  |
| Age(years), mean (SD)* | 10.40 (1.92) | 6.93 (2.53) | 9.30 (2.66) | <0.001 |
| Weight(kg), mean (SD) | 1.71 (0.33) | 1.60 (0.25) | 1.67 (0.31) | 0.4 |
| Length(cm), mean (SD) | 42.68 (2.38) | 41.64 (2.06) | 42.35 (2.31) | 0.2 |
| Histological Score, mean (SD) | 3.20 (1.03) | 0.86 (0.36) | 2.45 (1.41) | <0.001 |

\* Standard Deviation

#### 4.1.2 Preprocessing

Prior to statistical analysis, we preprocessed the metabolomic and lipidomic datasets through the following consecutive steps: (1) imputation based on half of the minimum value strategy, (2) we used the probabilistic quotient normalization method (Dieterle *et al*., 2006) for normalization, (3) we applied a log-transformation to make data more symmetric and homoscedastic, and (4) we accounted for batch effect using the surrogate variable analysis from the R package sva (version 3.42.0)(Leek *et al*., 2012) with the ComBat method (Johnson *et al*., 2007).

#### 4.1.3 Results

As we mention in the description of the study frame, our focus was on triglycerides, the most abundant lipid class in our dataset. Triglycerides consist of three fatty acid chains connected by a glycerol molecule. The heatmap of Figure 2 (a) clearly indicates that some correlation structure exists among the set of triglyceride levels, as shown by the correlation matrix. The length of each chain and its degree of unsaturation is determined by the number of carbons and double bonds present, respectively. By examining patterns across triglycerides or groups of triglycerides based on these intrinsic properties, we were able to gain insights into their profiles. Figure 2 (b) enables easy assessment of the effect of fish oil supplementation on different types of triglycerides. The t-statistics calculated for each triglyceride in Figure 2 would be a traditional method of examining the effect of fish oil supplementation.

**Figure 2:**
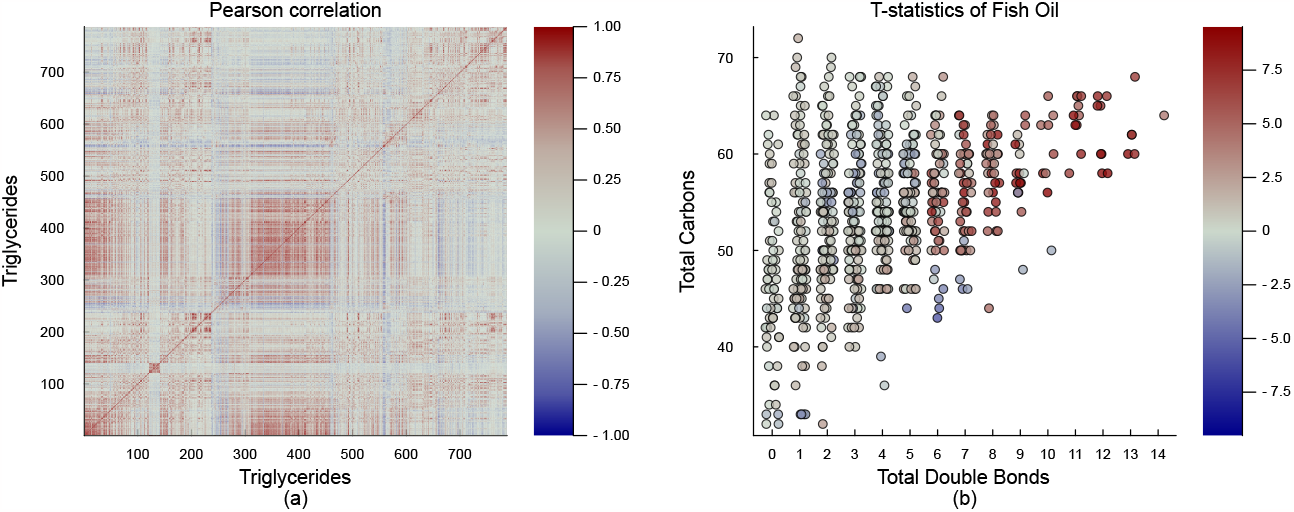
(a) Heatmap illustrating Pearson correlation coefficients among triglycerides. Shades of red denote strong positive correlations, grey signifies minimal or no correlation, and shades of blue represent strong negative correlations. (b) T-statistics representation for 787 triglycerides between individuals with and without fish oil supplementation. Each dot symbolizes a specific triglyceride. The color of the dot indicates the impact of fish oil supplementation: red dots signify an increase in the triglyceride level, while blue dots represent a decrease when compared to individuals without fish oil supplementation. The dot’s position relates to the total count of carbons and double bonds in that specific triglyceride.

The number of carbons in a fatty acid chain is closely related to the presence of double bonds. This correlation exists because as the chain gets longer, there are more opportunities for a double bond to form. When considering triglycerides as a whole, the effect of the number of carbons is intertwined with the number of double bonds. We can note in Figure 2 (b) that individuals who supplement with fish oil tend to have higher levels of triglycerides that contain a greater number of double bonds, typically more than 6 (i.e., more unsaturated fats). However, we were to perform an enrichment analysis of t-statistics using only the number of carbons. In that case, we might mistakenly conclude that triglycerides with more carbons are more abundant in people taking fish oil. Therefore, it is essential to consider both variables, the number of carbons and double bonds, to correctly interpret the results.

Matrix linear models are a powerful tool for disentangling the effects of different variables in complex datasets. Figure (3) illustrates the data structures of our matrix linear model as applied to the triglycerides dataset. In this example, the response matrix **Y** comprises the abundance of each triglyceride for all patients. Each row represents a separate patient sample, while each column corresponds to a specific triglyceride. The predictor matrix **X** contains information on the sample characteristics, such as whether patients received fish oil supplementation. The matrix **Z** provides information on the number of double bonds present in each triglyceride molecule. By incorporating data matrices **X** and **Z**, the matrix linear model enables us to assess the combined effects of fish oil supplementation and the degree of unsaturation on triglyceride levels. The matrix of coefficients **B** captures these relationships, allowing us to understand better the underlying biological mechanisms and potential interactions between the two factors.

**Figure 3:**
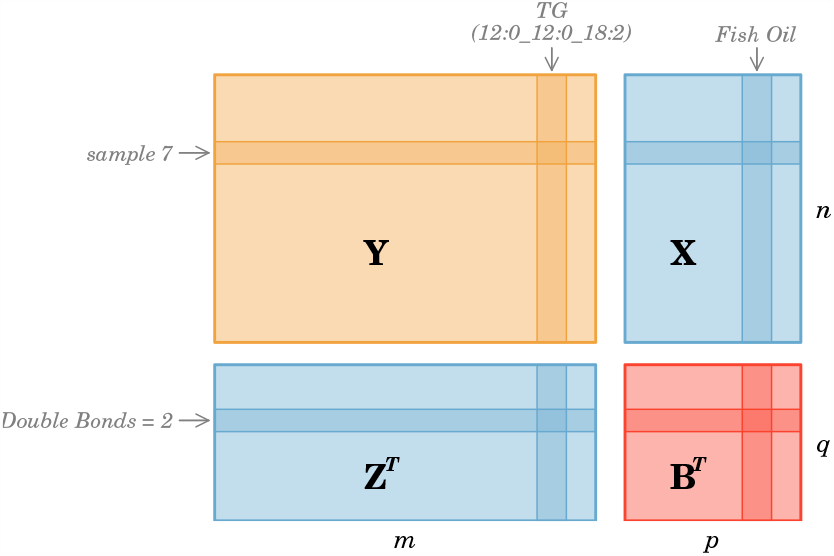
A visualization of the response (**Y** : *n*×*m*), covariates (**X** : *n*×*p*), measurement attributes (**Z** : *m*×*q*), and coefficients (**B** : *p*×*q*) matrices for a matrix linear model. The dimensions in the model correspond to *n* individuals, *m* measurements, *p* covariates, and *q* attributes; the matrix **B** is to be estimated.

In this particular example, the matrix linear model has been used to examine the relationship between fish oil supplementation and triglyceride levels while considering the number of carbons and double bonds in the triglycerides. By adjusting for both variables in the **Z** matrix, the analysis results in Figure 4 show that the effect of fish oil is mainly seen in triglycerides with 6 or more double bonds. This suggests that fish oil is particularly effective at increasing the levels of polyunsaturated triglycerides. Importantly, this effect is not driven by differences in chain length, which have been adjusted for in the analysis. This approach also highlights the importance of adjusting for confounding variables when analyzing complex datasets. Without adjusting for the number of double bonds, an unadjusted analysis would have suggested an effect of fish oil on both chain length and the number of double bonds. The adjusted analysis provides, therefore, a more comprehensive and informative view of the underlying biological mechanisms. Overall, this example demonstrates the potential power of matrix linear models in identifying and disentangling complex relationships between variables in biological systems.

**Figure 4:**
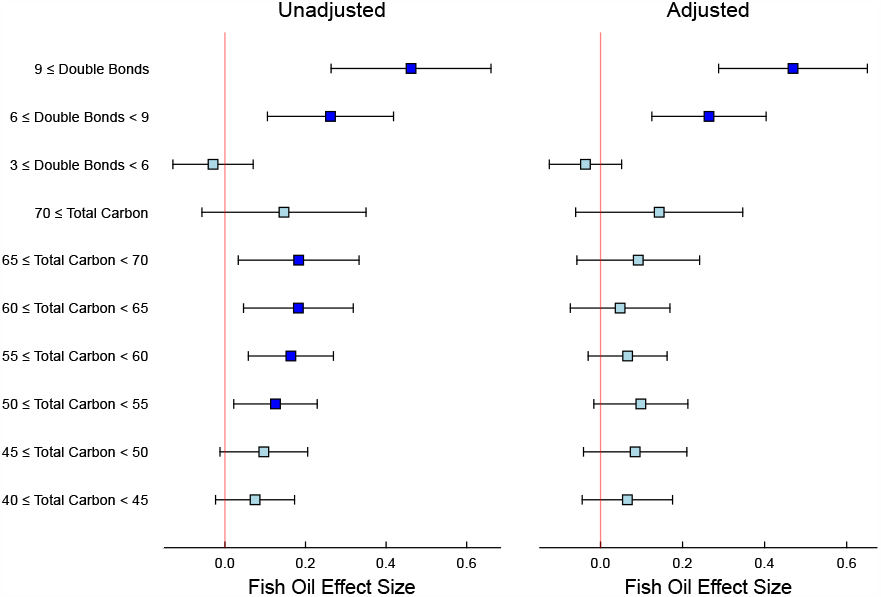
Statin study: Fish oil effect size in triglycerides. The left panel shows the unadjusted effects of the number of double bonds and number of carbons; the right panel shows the effects of the number of double bonds and the number of carbons adjusted for the other.

### 4.2 Pansteatatis Mozambique tilapia study

#### 4.2.1 Descriptive characteristics

In this study, 51 Mozambique tilapia (Oreochromis mossambicus) were captured from various locations around the inflow of Loskop Dam, South Africa, and underwent detailed analysis under an approved animal handling protocol. Immediately after capture, blood samples were collected and later analyzed using advanced liquid chromatography and high-resolution tandem mass spectrometry to characterize the plasma lipidome. Additionally, adipose tissue samples were histologically examined to assign disease severity scores, ranging from 0 (no disease) to 5 (severe disease). Fish with histological scores above 1 were categorized as diseased, and those with scores of 1 or less as healthy. However, due to missing data in the adipose tissue histological scores, our analysis was conducted on a subset of 44 tilapia instead of the initial 51. Table2 displays the descriptive statistics for the healthy and diseased tilapia groups.

#### 4.2.2 Preprocessing

The metabolomic datasets from the Metabolomics Workbench (see Data Analysis & Data Sources) had no missing data and underwent preprocessing that involved the following steps: (1) implementing a normalization approach using the probabilistic quotient normalization (Dieterle *et al*., 2006) method, and (2) performing a log2-transformation to increase data symmetry and homoscedasticity.

#### 4.2.3 Results

In the Mozambique tilapia study, we applied Matrix Linear Models to explore the influence of fish age on various lipid groups, considering different levels of annotation. The covariates included health status, length, and weight. We used a few different versions of the **Z** matrix as outlined below. Each version helps us answer slightly different questions.

Initially, we employed superclass information, identifying four primary groups: sterol lipids, sphingolipids, glycerophospholipids, and glycerolipids. As the confidence plot of the Figure 5(a) demonstrates, older fish exhibit a lower abundance of glycerolipids and a higher abundance of glycerophospholipids and sterol lipids. Upon examining subclasses for more granular annotations, we observed that with increasing fish age, triglyceride levels decrease while phospholipid levels, particularly phosphatidylcholines and phosphatidylethanolamines, increase (Figure 5(b)).

**Figure 5:**
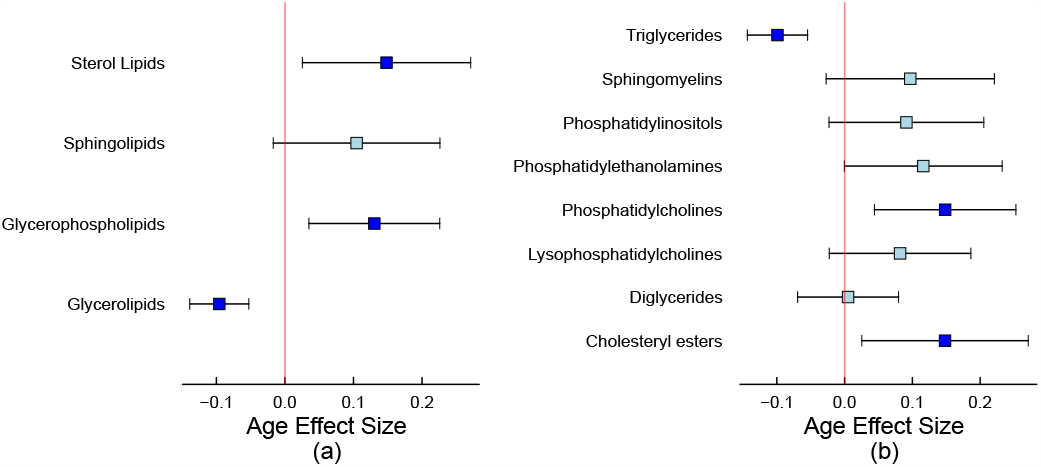
Pansteatitis study: Confidence plot showing the age size effect on the lipid (a) superclasses and (b) subclasses.

Lastly, we investigated patterns across triglyceride groups based on carbon chain length and degree of unsaturation. Adjusting for both variables in the Z matrix, our analysis in Figure 6 reveals that the age effect is primarily observed in triglycerides with fewer than 65 carbons, without being driven by differences in unsaturation levels. This approach underscores the necessity of accounting for confounding variables when analyzing complex datasets. Without adjusting for both features, an unadjusted analysis erroneously suggested an age effect on triglycerides driven by the number of double bonds and carbon chain length.

**Figure 6:**
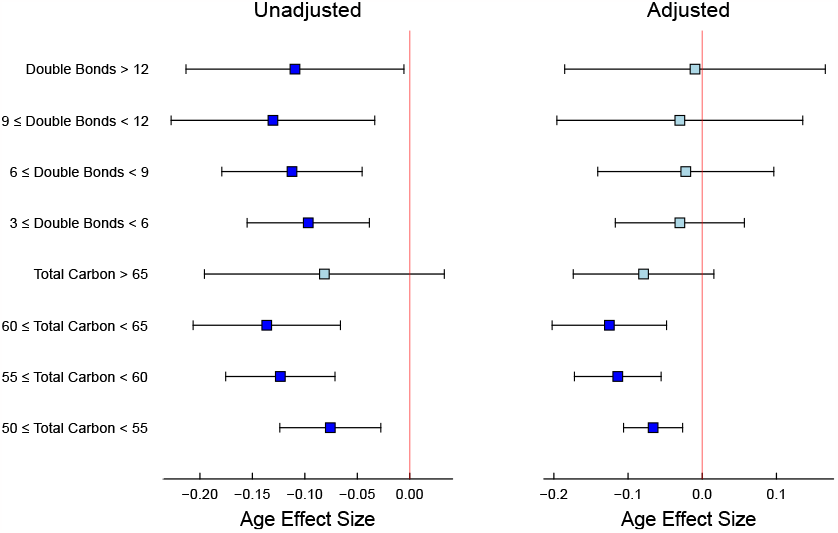
Pansteatitis study: Age effect size in triglycerides. The left panel shows the unadjusted effects of the number of double bonds and number of carbons; the right panel shows the effects of the number of double bonds and the number of carbons adjusted for the other.

### 4.3 COPD-SPIROMICS study

#### 4.3.1 Descriptive characteristics

The categorization of metabolites into “Super Class” and “Sub Class” was carried out by Metabolon (Durham, NC, USA). The COPDGene cohort showed that males were significantly older, had more smoking packyears (although fewer current smokers), had a higher percentage of COPD cases, and had a higher mean percentage of emphysema than females (Gillenwater *et al*., 2021). A similar pattern of sex-specific differences was observed for smoking pack-years and COPD cases in the SPIROMICS cohort. The COPDGene cohort generally comprised older individuals with a lower proportion of African-American subjects, fewer current smokers, less smoke exposure measured by cigarette pack years, and a lower emphysema percentage than the SPIROMICS cohort.

#### 4.3.2 Preprocessing

To preprocess the metabolomic datasets, we followed a series of steps, which included: (1) applying a KNN imputation using the nearest neighbor averaging method from the R package “impute” (Hastie *et al*., 2023), (2) performing a normalization using the probabilistic quotient normalization method (Dieterle *et al*., 2006), and (3) applying a log2-transformation to make the data more symmetric and homoscedastic.

#### 4.3.3 Results

In the context of the COPD study, we investigated the impact of sex on metabolites using Matrix Linear models with three different Z matrices, each containing information on the superclass, subclass, and modules obtained from Weighted Gene Co-expression Network Analysis (WGCNA) clustering algorithms (Gillenwater *et al*., 2021). Gillenwater et al. (2021) used WGCNA to cluster metabolite abundances into modules of correlated metabolites and identified 11 co-varying metabolite modules, mainly separated based on metabolite subclass (see Table 4). This approach allowed us to explore subnetworks differentially dysregulated within COPD subjects by sex and identify sex-specific biomarkers of those subnetworks. We performed Matrix Linear Models with metabolites as the outcome and COPD phenotype as the predictor, along with the following covariates: sex, age, race, BMI, current smoking status, smoking pack-years, clinical center, percent emphysema and FEV1pp (Forced Expiratory Volume percent predicted).

At the superclass level, we observed that the Xenobiotics, Nucleotide, Lipid, Energy, and Amino Acid superclasses showed substantial differences between sexes in the COPDGene study (Figure 7). Females had higher metabolite abundances in the Lipid and Energy superclass, while males had higher quantities in the Xenobiotic, Nucleotide, and Amino Acid superclasses. The MLM framework allows for comparative analysis between models derived from the SPIROMICS and COPDgene datasets. This approach centers on assessing the differences and averages of select regression coefficients contingent on shared variables or factors and calculating associated standard errors and confidence intervals. The COPDGene and SPIROMICS studies confirmed that females had higher levels of Lipid and Energy metabolites and lower levels of Nucleotide and Amino Acid metabolites than males, with two exceptions: Xenobiotics superclass was only significant in COPDGene, and males had substantially higher abundances in the Peptide superclass.

**Figure 7:**
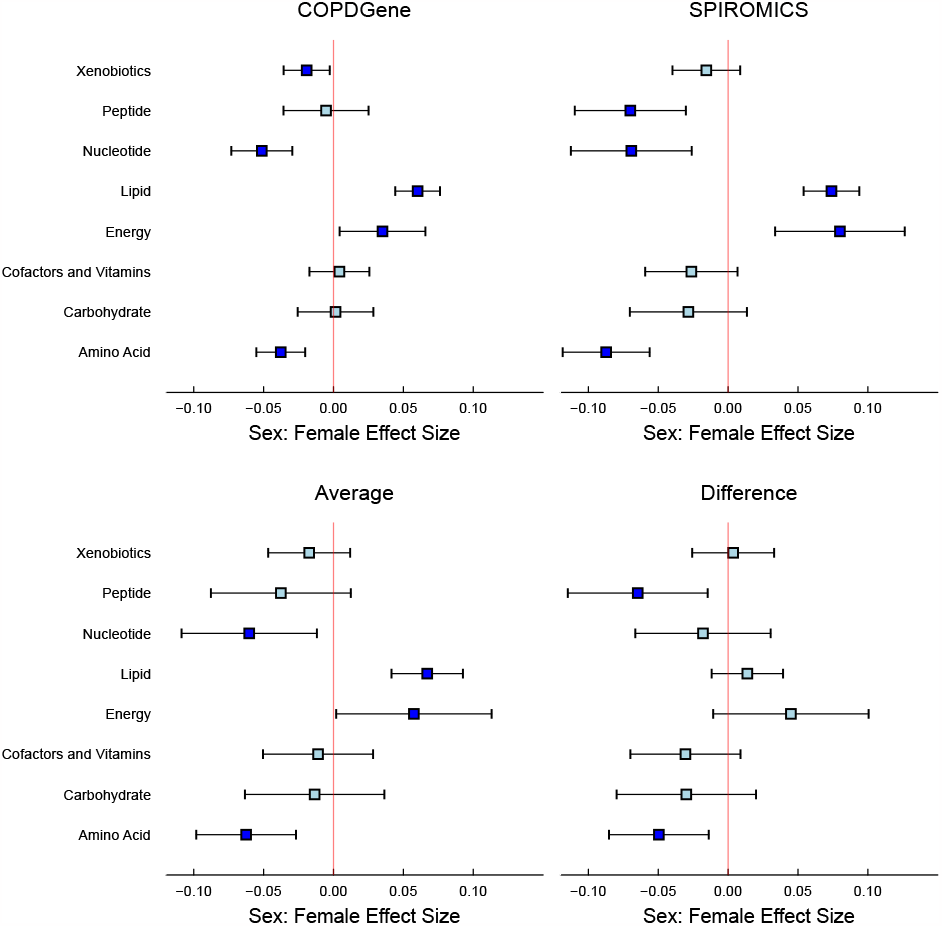
Comparison of COPDGene and SPIROMICS studies: Confidence plots comparing the similarities and differences in the female sex effect between both cohorts by metabolite class.

The second Z matrix design allowed us to magnify information by examining subclasses’ annotations, enabling the identification of significant subclasses. We observed substantial differences between females and males in the Lipids, Amino Acids, and Nucleotides subclasses (Figure 8). The most notable differences between sexes were that females had higher Sphingomyelin metabolite levels, while males had higher Androgenic Steroid levels (Table 3).

**Table 3:** Sorted Subclasses by average effect size of COPDGene and SPIROMICS studies, with a significance level of FDR < 0.05.

| ID | name | Effect Size | Conf. Interval | FDR |
| --- | --- | --- | --- | --- |
| LIP01 | Androgenic Steroids | -0.28 | [-0.36, -0.21] | 0.000 |
| AMI08 | Leucine, Isoleucine and Valine Metabolism | -0.13 | [-0.18, -0.08] | 0.000 |
| AMI11 | Phenylalanine Metabolism | -0.12 | [-0.18, -0.06] | 0.000 |
| LIP15 | Fatty Acid Metabolism (Acyl Carnitine, Polyunsaturated) | -0.11 | [-0.22, -0.01] | 0.030 |
| NUC01 | Purine Metabolism, (Hypo)Xanthine/Inosine containing | -0.10 | [-0.16, -0.05] | 0.001 |
| AMI07 | Histidine Metabolism | -0.08 | [-0.13, -0.02] | 0.012 |
| AMI13 | Tryptophan Metabolism | -0.08 | [-0.13, -0.04] | 0.001 |
| AMI03 | Glutamate Metabolism | -0.07 | [-0.12, -0.02] | 0.008 |
| NUC07 | Pyrimidine Metabolism, Uracil containing | -0.07 | [-0.13, -0.02] | 0.012 |
| LIP27 | Fatty Acid, Monohydroxy | 0.06 | [0.00, 0.12] | 0.040 |
| LIP36 | Lysophospholipid | 0.10 | [0.02, 0.17] | 0.012 |
| LIP08 | Endocannabinoid | 0.13 | [0.04, 0.22] | 0.008 |
| LIP42 | Phosphatidylethanolamine (PE) | 0.13 | [0.02, 0.23] | 0.016 |
| LIP38 | Medium Chain Fatty Acid | 0.14 | [0.07, 0.22] | 0.000 |
| LIP35 | Long Chain Saturated Fatty Acid | 0.15 | [0.04, 0.26] | 0.011 |
| LIP41 | Phosphatidylcholine (PC) | 0.15 | [0.08, 0.22] | 0.000 |
| LIP44 | Phosphatidylinositol (PI) | 0.15 | [0.05, 0.25] | 0.006 |
| LIP29 | Hexosylceramides (HCER) | 0.17 | [0.08, 0.26] | 0.001 |
| LIP33 | Long Chain Monounsaturated Fatty Acid | 0.18 | [0.08, 0.28] | 0.001 |
| LIP34 | Long Chain Polyunsaturated Fatty Acid (n3 and n6) | 0.18 | [0.08, 0.28] | 0.001 |
| LIP07 | Dihydrosphingomyelins | 0.19 | [0.09, 0.29] | 0.000 |
| LIP54 | Sphingomyelins | 0.31 | [0.22, 0.39] | 0.000 |

**Table 4:** Metabolite Classes by modules determined using WCGNA.

| Module | Metabolite Classes |
| --- | --- |
| blue | Acyl Carnitines, Fatty Acids (Dicarboxylate, Monohydroxy, Long chain, Medium chain), Endocannabinoids, Nucleotides |
| red | Ceramides, Sphingomyelins |
| turquoise | Xenobiotics, Amino Acids (Tryptophan metabolism, Glutamate metabolism, Histidine metabolism, Branched Chain Amino Acids, Glycine, Serine and Threonine Metabolism, Methionine, Cysteine, SAM and Taurine Metabolism, Polyamine Metabolism, Urea cycle; Arginine and Proline Metabolism), TCA cycle metabolites |
| brown | Amino Acids (Gamma-glutamyl Amino Acid, Glutamate Metabolism, Branched Chain Amino Acids, Urea cycle; Arginine and Proline Metabolism, Lysine Metabolism, Methionine, Cysteine, SAM and Taurine Metabolism, Phenylalanine Metabolism), Bile Acids, Acyl Cholines, Lysophospholipids |
| yellow | Xenobiotics (Benzoates, Xanthines, Nutritional) |
| green | Lysophospholipids, Phosphatidylcholines (PC), Phosphatidylinositols (PI), Plasmalogens |
| magenta | Steroids (Androgenic, Pregnenolone, Corticosteroids, Progestin) |
| black | Diacylglycerols, Phosphatidylethanolamines (PE), Acyl Carnitines |
| greenyellow | Cofactors and Vitamins |
| purple | Acetylated peptides, Xenobiotics (Benzoates), Secondary Bile Acids |
| pink | Xenobiotics (Chemicals), Dipeptides, Hemoglobin and Porphyrin Metabolites |

**Figure 8:**
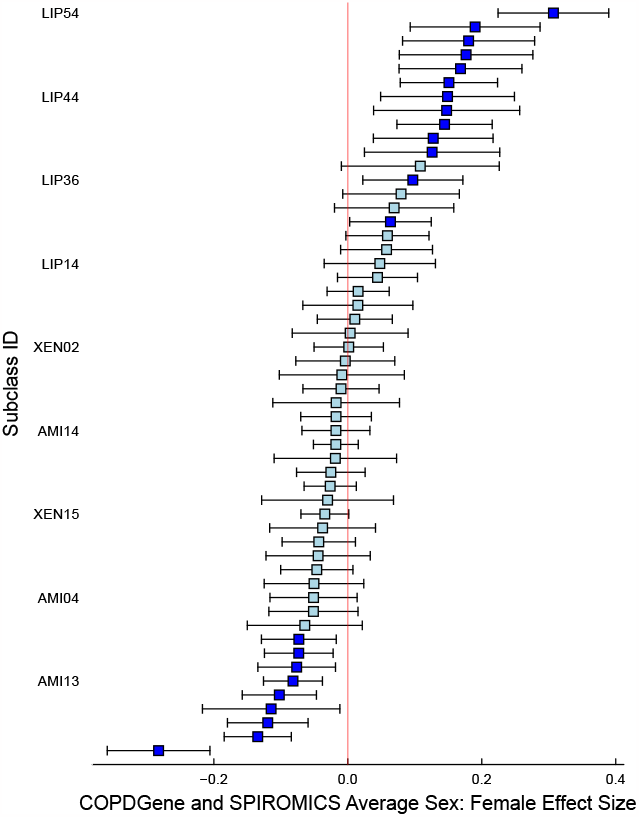
Sorted average sex effect by metabolite subclass: Confidence interval plots showing the sorted average effect in the two studies by subclasses present in both the COPDGene and SPIROMICS cohorts.

MLM results based on the WGCNA modules aligned closely with the findings of Gillenwater et al. (2021). In the COPDGene cohort, sex-specific differences were significant in 7 out of 11 identified modules. The red and magenta modules had the most pronounced differences, with females having higher metabolite abundances in the red module and males having higher abundances in the magenta module (Figure 9). Similarly, the blue and green modules showed higher abundances in females, while the turquoise, pink, and brown modules had higher abundances in males. SPIROMICS showed a similar trend, with 7 out of 11 modules significantly differing by sex. However, there were two dissimilarities with the COPDGene results: the pink module was insignificant, and the purple module had significantly higher levels of metabolites in females. On average, the effect of both cohorts indicated that only five modules remained significant: blue, red, turquoise, green, and magenta.

**Figure 9:**
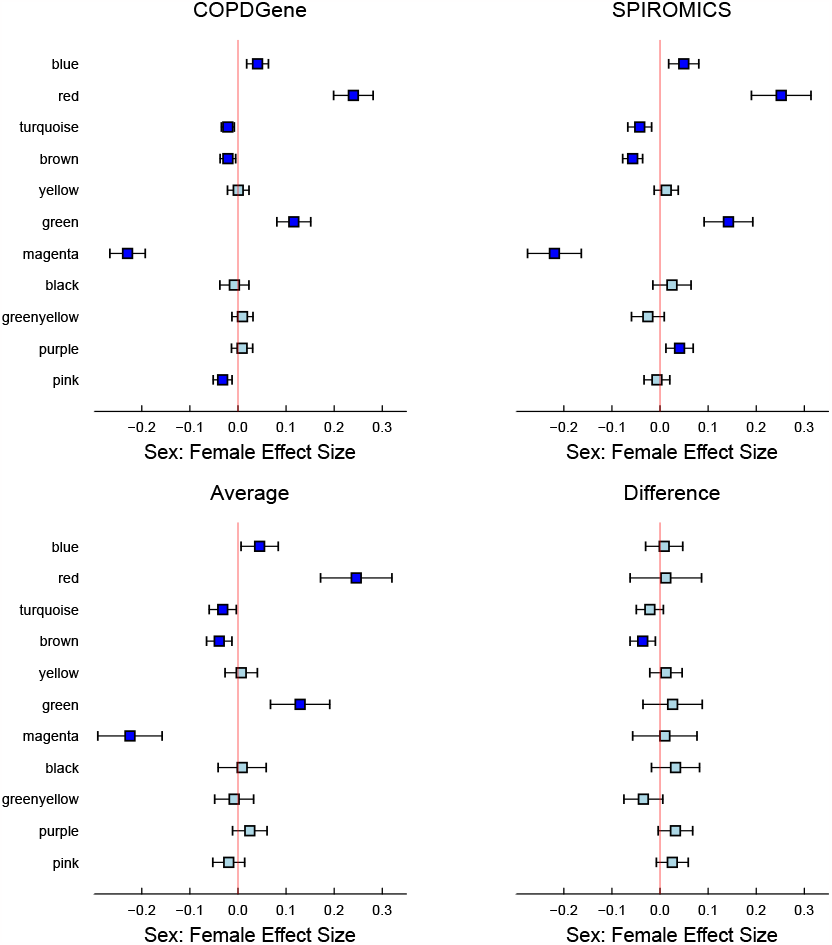
Comparison of COPDGene and SPIROMICS studies: Confidence plots comparing the similarities and differences in the female sex effect between both cohorts by modules determined using WCGNA in COPDGene and SPIROMICS studies. 6 of the 11 modules showed a significant effect size associated with sex:female in both cohorts, indicating a strong and consistent association. Except for the brown module, which showed a distinct variation in effect size, the remaining modules presented statistically comparable results across both cohorts. The direction of the sex:female effect was uniform across both cohorts.

## 5 Discussion

We analyzed three distinct datasets using matrix linear models to uncover complex relationships between variables in biological systems. The three examples provided insights into the effect of fish oil supplementation on triglycerides (SAMS study), the impact of age on lipid profiles in fish (Pansteatitis study), and sex-specific differences in the metabolite profiles of COPD patients (COPD-SPIROMICS study). A key feature of our analytic approach is to explicitly use the metabolite annotations using a bilinear model. This enables us to aggregate signals in metabolites sharing one or more annotations, and can detect associations that may be individually undetectable.

In the SAMS study, we examined the effect of fish oil supplementation on triglyceride levels, considering the number of carbons and double bonds in triglyceride molecules. Our analysis demonstrated that fish oil primarily increased the levels of polyunsaturated triglycerides with six or more double bonds. This finding suggests that fish oil supplementation is particularly effective at increasing levels of polyunsaturated fats, which are known to benefit cardiovascular health. Moreover, this example highlights the importance of examining two or more metabolite covariates jointly to separate the effects of each. This is a phenomenon similar to confounding in epidemiological studies, where the effect of one variable (number of double bonds) may be masked by the effect of another (number of carbon atoms).

In the Pansteatitis study, we focused on the impact of age on lipid profiles in fish, revealing that older fish exhibit a decrease in triglyceride levels and an increase in phospholipid levels, particularly phosphatidylcholines, and phosphatidylethanolamines. This analysis also demonstrated the importance of adjusting for confounding variables, such as carbon chain length and degree of unsaturation, to obtain accurate insights into the age effect on lipid profiles. This approach may provide valuable information for understanding the aging process and its impact on lipid metabolism in aquatic organisms.

In the COPD-SPIROMICS study, we investigated sex-specific differences in the metabolite profiles of COPD patients. Our analysis revealed significant differences between males and females in various metabolite superclasses, subclasses, and co-varying metabolite modules. Notably, females had higher lipid and energy metabolite levels, while males had higher nucleotide, amino acid, and xenobiotic metabolite levels. These findings suggest that sex-specific differences in metabolite profiles might be relevant to the pathophysiology and treatment of COPD. In this study, our framework allowed us to estimate the average effect of a group of metabolites across the two studies and also show where they differ in an interpretable manner. Utilizing the clusters defined in the original methodology yields similar results, demonstrating the robustness of our approach. For researchers who prefer not to use cluster-based analysis, our method allows for directly identifying associated annotations or classes, offering a more tailored analytical perspective. Our approach enables the summarization and contrasting of two studies with the added confidence intervals, facilitating a more nuanced understanding of the data. The Sphingomyelins subclass exhibited the most substantial effect size (FDR < 0.001) among the metabolites analyzed (Table 3). This finding is congruent with the red module’s significance (Figure 9), which encompasses Sphingomyelins among its metabolite classes and shows a strong effect size. Our analysis revealed that females have significantly lower levels of Androgenic Steroids compared to males, which aligns with the results from the magenta module, includes the subclass Androgenic Steroids. This coherence between subclass metabolite levels and module composition strengthens the reliability of the observed sex-specific differences in metabolomic profiles. Gillenwater et al. (2021)’s research corroborates that metabolomic profiles exhibit pronounced sex-based differences, particularly within distinct modules of sphingolipids and steroids. Their methodology involved a three-step process: first clustering the data, then associating these clusters with sex, and finally determining the contributing annotations to the sex-associated clusters. In contrast, our approach streamlines this process into a single step by employing the Z matrix of the matrix linear model (MLM) to directly analyze the subclass annotations, thereby simplifying and potentially enhancing the efficiency of the analysis.

Our results demonstrate the power and utility of matrix linear models in disentangling complex relationships between variables in biological systems. A novel feature of the MLM is in the introduction of the metabolite characteristic matrix (aka the *Z* matrix). This innovative approach enables researchers to identify and assess interesting subgroups of metabolites based on their attributes and known groupings, such as pathways, class, subclass, or quantitative molecular features. The *Z* matrix serves as a window into the metabolite features, allowing for treating metabolites as a population and exploring compelling subgroups. In terms of data processing, MLM streamlines the analysis by investigating analyzing the metabolites in a single step, as opposed to the conventional two-step approach, which is time-consuming and less efficient. The MLM is estimated using a computationally efficient algorithm based on matrix multiplications, that is faster than analyzing the metabolites one by one. It offers a simple and flexible framework that maintains interpretability while efficiently processing complex datasets. By incorporating external information typically ignored in standard high-throughput data analyses, MLM provides a flexible tool that can enhance the overall understanding of complex biological systems.

Despite these advantages, the MLMs should be viewed as another tool in the data analyst’s toolbox which should be deployed as the situation demands. For example, when each metabolite is of individual interest, then they should be analyzed individually (by using the identity matrix as the *Z* matrix, as this is faster than looping over individual regressions). There is also the implicit assumption that all metabolites sharing a characteristic have similar effects in the same direction. If it is suspected that the effects may be in different directions, a different approach such as a kernel-based approach might be superior. Our current approach assumes that the variance does not depend on the mean. For binary or count data, the variance and the mean might be linked, and for such data, generalizations of our approach need to be developed. Since we are using a squared error loss function for estimation, our approach may be sensitive to outliers. The data analyst is advised to remove such outliers or suitably process them by making data transformations. In the future, we expect to develop approaches that use more robust loss functions.

In conclusion, the matrix linear model presents a significant advancement in the field of metabolomics data analysis. By offering a flexible, efficient, and interpretable framework for incorporating external information and exploring complex relationships between variables, MLM has the potential to greatly improve our understanding of biological systems and inform the development of targeted interventions for various health conditions.

## Software

Our implementation of MLMs in Julia. the `MatrixLM` package is available at https://github.com/senresearch/MatrixLM.jl. The `MetabolomicsWorkbenchAPI` package is available at https://github.com/senresearch/MetabolomicsWorkbenchAPI.jl.

## Data Analysis & Data Sources

Interactive notebooks in Pluto are available from our Github site: https://github.com/senresearch/mlm-metabolomics-supplement.

Pansteatatis Mozambique Tilapia data are available at the NIH Common Fund’s National Metabolomics Data Repository (NMDR) website, the Metabolomics Workbench, https://www.metabolomicsworkbench.org, where it has been assigned Project ID PR000705. The data can be accessed directly via its Project DOI: 10.21228/M8JH5X This work is supported by Metabolomics Workbench/National Metabolomics Data Repository (NMDR) (grant# U2C-DK119886), Common Fund Data Ecosystem (CFDE) (grant# 3OT2OD030544) and Metabolomics Consortium Coordinating Center (M3C) (grant# 1U2C-DK119889).

COPDGene and SPIROMICS data are available at the NIH Common Fund’s National Metabolomics Data Repository (NMDR) website, the Metabolomics Workbench, https://www.metabolomicsworkbench.org, where it has been assigned Project ID PR001048. The data can be accessed directly via its Project DOI: 10.21228/M87D6G This work is supported by Metabolomics Workbench/National Metabolomics Data Repository (NMDR) (grant# U2C-DK119886), Common Fund Data Ecosystem (CFDE) (grant# 3OT2OD030544) and Metabolomics Consortium Coordinating Center (M3C) (grant# 1U2C-DK119889).

## Acknowledgements

We acknowledge the support of NIH grants R21AR0704018 (MBE, GF, SS), R01GM123489 (GF, SS), P30DA044223 (KK,SS).

## Author contributions

- Data analysis: GF, CZ, SS
- Statistical methods: GF, KK, SS
- Biological interpretation: TG, MBE, KK
- First draft: GF
- Review and editing: GF, HYC, KK, TG, MBE, SS

